# The commensal protist *Tritrichomonas musculus* exhibits a dynamic life cycle that induces extensive remodeling of the gut microbiota

**DOI:** 10.1101/2023.03.06.528774

**Authors:** Ana Popovic, Eric Yixiao Cao, Joanna Han, Nirvana Nursimulu, Eliza V.C. Alves-Ferreira, Kyle Burrows, Andrea Kennard, Noor Alsmadi, Michael E. Grigg, Arthur Mortha, John Parkinson

## Abstract

Commensal protists and gut bacterial communities exhibit complex relationships, mediated at least in part through host immunity. To improve our understanding of this tripartite interplay, we investigated community and functional dynamics between the murine protist *Tritrichomonas musculus* (*T. mu*) and intestinal bacteria in healthy and B cell-deficient mice. We identified dramatic, protist-driven remodeling of resident microbiome growth and activities, in parallel with *T. mu* functional changes, accelerated in the absence of B cells. Metatranscriptomic data revealed nutrient-based competition between bacteria and the protist. Single cell transcriptomics identified distinct *T. mu* life stages, providing new evidence for trichomonad sexual replication and the formation of pseudocysts. Unique cell states were validated *in situ* through microscopy and flow cytometry. Our results reveal complex microbial dynamics during the establishment of a commensal protist in the gut, and provide valuable datasets to drive future mechanistic studies.

## Introduction

Research into host-associated microbiomes has established the critical role of gut bacteria in health and disease^1,2^. Emerging data supports an appreciable presence of gut-dwelling protists with host immunomodulatory capabilities and influence over resident bacterial communities, suggesting these microbes are essential players in organizing the gut environment^3–7^. Studies in this discipline are in their infancies however and lack understanding of both the ability of protists to colonize and persist and protist:bacterial cross-talk in the context of host immunity.

The protist *Tritrichomonas musculus* (*T. mu*) is a common resident of the murine gut, and a relative of prevalent human trichomonads with links to colitis, irritable bowel syndrome and colon cancer^5,8–12^. *T. mu* establishes chronic asymptomatic infections, resulting in elevated baseline intestinal immune activation and structural remodelling of the intestinal epithelium^6,13–15^. Protist-produced succinate furthermore stimulates a Th2-based immune response that restricts infection by enteric parasites^14–16^. Although the sustained inflammation associated with *T. mu* confers protection to mice against pathogenic bacteria, it can increase susceptibility to T-cell-driven colitis and tumorigenesis, suggesting *T. mu* behaves as a pathobiont. Evidence shows its presence influences gut bacterial composition, and that conversely *T. mu* engraftment may be modulated by bacterial taxa such as *Bifidobacterium* spp. through unknown mechanisms^17^. These microbial interactions may in part be driven through host immunoglobulins (Ig) A and M, antibodies secreted by B cells and elevated in response to *T. mu*^6,15^. Both bacteria and protists have known effects on B cell antibody responses, but it is not well understood how host B cells modulate cross-kingdom microbial interactions^6,18,19^.

Here we track *T .mu* activities during the first 28 days of gut colonization, together with its interactions with resident bacteria in the context of a healthy WT host and B cell-deficient (*muMT^-/-^*) mice through 16S sequence surveys and microbial transcriptomics. We show that *T. mu* expansion is accelerated in the absence of host B cells. Its colonization dramatically remodels bacterial composition and function, concurrent with shifting protist metabolism and virulence factor expression - changes accelerated in the absence of host B cells. We provide detailed transcriptomic analyses suggesting cross-kingdom competition for key dietary nutrients. Finally, we conduct single-cell level characterization of the *T. mu* life cycle and validate cell stages *in situ* through fluorescence-based labelling of stage-specific transcripts. Our data reveal B cell-modulated co-adaptation between resident bacteria and a newly colonizing protist, and provide valuable datasets to drive future mechanistic studies.

## Materials and Methods

### Mice

C57Bl/6J and B6.129S2-*Ighm^tm1Cgn^*/J (*muMt*^-/-^) female mice (Jackson Laboratory, Bar Harbor, ME, USA) were maintained in specific pathogen free (SPF) conditions at the University of Toronto Division of Comparative Medicine. Experiments were conducted at 6-8W of age, using age and sex matched controls. For single-cell protist sequencing, 12W germ-free (GF) female C57Bl/6 mice (Taconic Biosciences, Germantown, NY, USA) were housed in the GF facility until the day of analysis, when they were transported to SPF conditions. Mice conventionalized with a microbiome received a bacterial suspension through oral gavage derived from two C56Bl/6J fecal pellets in PBS, and were maintained under SPF conditions for 4 weeks prior to protist colonization. Animals were housed in a closed caging system and provided with an irradiated chow diet (Envigo Teklad 2918), non-acidified water (reverse-osmosis and UV-sterilized) with a 12hr light/dark cycle. Following *T. mu* colonization, animals were housed in separate cages. Animal experiments were approved by the Local Animal Care Committee (LACC) at the Faculty of Medicine, University of Toronto (animal use protocol 20012400 to AM).

### Protist colonization

Purification of *T. mu* was performed as previously described^6^. Briefly, cecal contents of a colonized C57Bl/6J mouse were filtered through a 70µm cell strainer, washed with PBS, and protists were collected from the interphase after density centrifugation through 40% Percoll overlaid on 80% Percoll. Cells were sorted into PBS on a BD Influx Cell Sorter (BD Biosciences, Franklin Lakes, NJ, USA) using the 100µm nozzle at 27psi at 4°C, with >99% purity. Mice were orally gavaged immediately after with two million *T. mu* cells. Protists were quantified using a hemocytometer.

### Genome sequencing and annotation

Genomic DNA was extracted from sorted protists using the MagAttract HMW DNA Kit (QIAGEN, Hilden, Germany), and sequenced using PacBio Sequel technology on two SMRT cells at the McMaster University Farncombe Metagenomics Facility (Hamilton, Canada). Reads were error corrected using CANU v1.8^20^, assembled into contigs using Flye v.2.4.1^21^, and subsequently polished with 8.6 million 300 bp paired-end Illumina MiSeq reads sequenced at the National Institute of Allergy and Infectious Diseases (Bethesda, Maryland) using BWA^22^ and Pilon v.1.23^23^. Annotation was carried out with the Maker v2.31 pipeline^24^ using SNAP v.2013-11-29^25^ and Augustus v.3.3.1^26^ for gene model prediction. Functional annotation was performed with InterProScan v.5.30-69.0^27^, HmmerWeb v.2.41.2^28^ and Architect^29^. Genes encoding adhesins, meiosis and cell cycle-related proteins were identified based on sequence homology with *T. vaginalis* proteins retrieved from the TrichDB database^30–32^. Detailed methods are provided in supplementary information. The genome assembly is accessible at: https://github.com/ParkinsonLab/Tritrichomonas-murine-microbiome-interactions/draft-genome-assembly.

### Phylogenetic analysis

Parabasalid ribosomal internal transcribed spacer (ITS) sequences were downloaded from GenBank^33^ (Table S12) and compared to ITS in the *T. mu* genome assembly. Multiple sequence alignments were generated using MUSCLE, and an unrooted phylogeny was constructed using the Maximum-likelihood method with a Tamura-Nei model in the MEGA-X software^34–36^. Bootstrap values were generated using 1000 replicates.

### scRNA-Seq analysis

A GF and conventionalized mouse were colonized with *T. mu* through oral gavage, as described above. Four weeks post colonization, protists were purified from caecal contents and immediately transferred on ice to the Princess Margaret Genomics Centre (Toronto, Canada) for STAMP library preparation using Drop-seq technology and sequenced on a NextSeq 500 (Illumina, San Diego, CA, USA)^37^. Reads were processed using Drop-seq Tools v.1.13 and aligned to the protist genome assembly using STAR v. 2.5.3a^38,39^. Three thousand protists per mouse (minimum 200 genes, 500 transcripts) were analyzed in Seurat v4 and the two transcriptomes were aligned using the FindIntegrationAnchors feature^40,41^. Cells were grouped using graph-based clustering (0.8 resolution, 18 principal components) and visualized via UMAP^42^. Differentially expressed (DE) genes were identified using the FindAllMarkers function, and functional enrichments were determined based on overrepresentation of pathway enzymes as defined by KEGG using the hypergeometric test, or GO terms using the topGO package and the Fisher’s Exact test^43,44^. Enrichments of custom-defined gene sets (meiosis, G1/S and G2 phase genes) were scored with the AddModuleScore function, and evaluated using two-sided Wilcoxon rank-sum tests. Benjamini-Hochberg correction was applied for multiple testing^45^. Heatmaps were generated using pheatmap 1.0.12 and Ward.D2 clustering^46^.

### 16S microbiome profiling and metatranscriptomics

Groups of four WT or *muMt*^-/-^ 6W female mice were infected with protists for microbial composition and gene expression profiling. A schematic of the experiment is presented in Fig. 1b.

**Figure 1.**
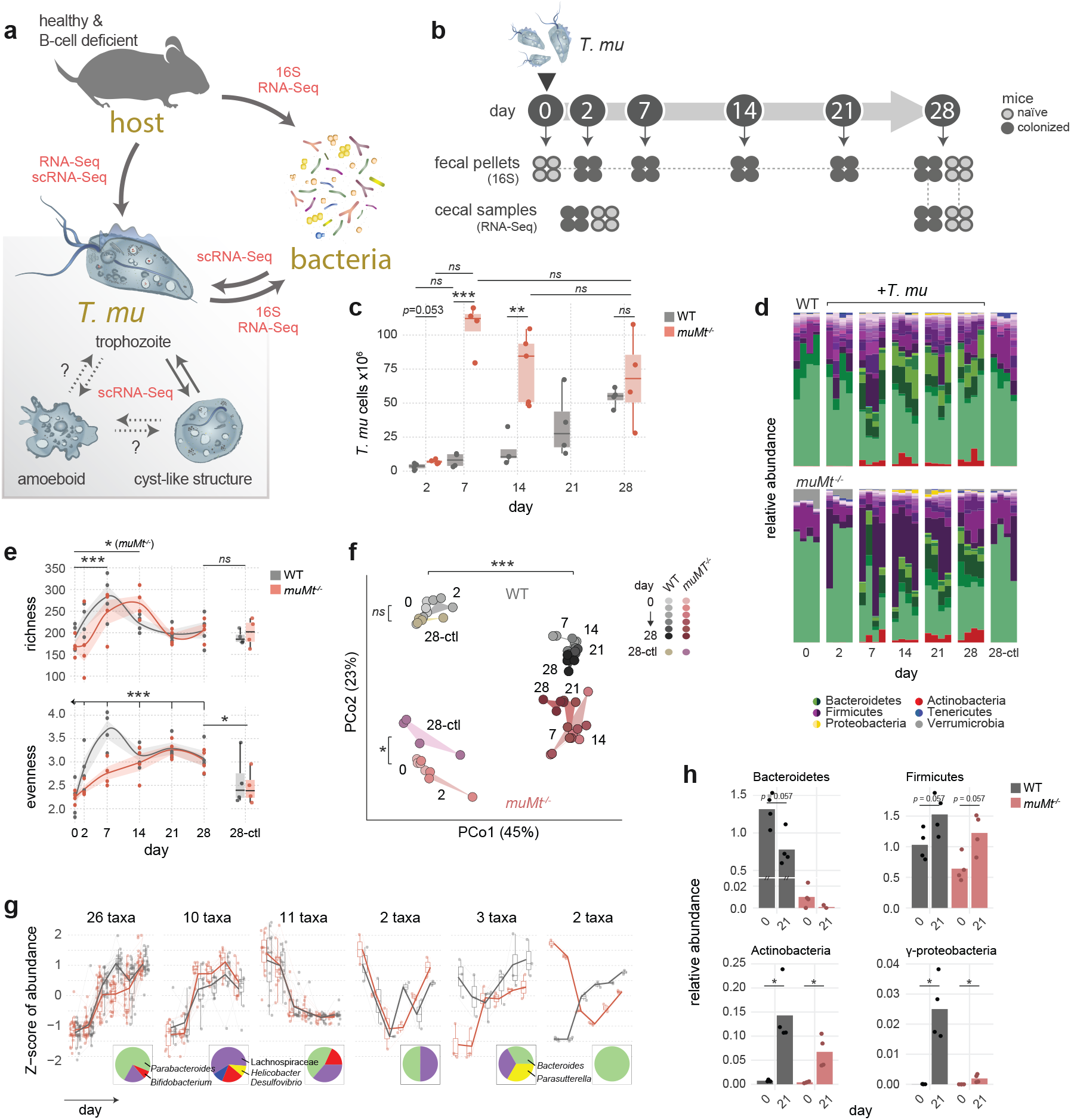
*T. mu* colonization drives bacterial diversification. **a**, Schematic of microbial interactions explored in the present study using 16S sequencing, metatranscriptomics (RNA-Seq) and/or single-cell transcriptomics (scRNA-Seq). **b**, Experimental design. Groups of four WT and *muMT^-/-^* mice were colonized with *T. mu*. Fecal pellets and caecal samples were collected at indicated timepoints for 16S and metatranscriptomic profiling. **c**, Expansion of *T. mu* cells in WT and *muMt*^-/-^ mouse caeca over 28 days of colonization (n=4 or 5 per group). Differences were tested using a two-sided t-test. **d**, Relative abundance of bacterial taxa determined through 16S sequencing of mouse fecal pellets from two groups of WT and *muMt*^-/-^ mice sampled over 28 days of colonization, and two groups of uninfected controls sampled on day 28 (n=4 per group). Colours represent genus-level abundances within indicated phylum colour groups. **e**, Bacterial richness (observed OTUs) and evenness (Shannon diversity) over the course of *T. mu* colonization. Differences between timepoints were evaluated using LME, and between colonized and naïve mice at day 28 using two-way ANOVA. **f**, Principal coordinate analysis of Bray-Curtis dissimilarities between samples. Significance was tested using permutational variance in Adonis. **g**, Patterns of bacterial abundance changes during *T. mu* colonization in WT (grey) and *muMt*^-/-^ (red) mice. Pie charts show the taxonomic make up of each cluster. **h**, Phylum-level bacterial abundance before (day 0) or 21 days after protist colonization, determined through qPCR using clade-specific primers. Abundances were normalized to Ct values obtained with universal 16S rRNA probes. Bars show means. Significance was evaluated using the Wilcoxon rank-sum test: \**p* < 0.05, ** < 0.01, *ns* not significant.

#### 16S profiling

Mouse fecal pellets were collected from the same group of WT or *muMT^-/-^* mice at days 0, 2, 7, 14, 21 and 28, or uninfected control mice at day 28, and stored at -80°C. DNA was extracted using the DNeasy PowerSoil Kit (QIAGEN, Hilden, Germany) according to manufacturer instructions, with a 5 min bead beating step using the FastPrep-24 homogenizer (MP Biomedicals, Santa Ana, CA, USA) at 5M/s. 16S rRNA amplicons were generated using B969F and BA1406R primers and paired-end sequenced (250 bp) with MiSeq v2 chemistry (Illumina, San Diego, CA, USA) at the Integrated Microbiome Resource facility at Dalhousie University (Halifax, Canada)^47^. Reads were pre-processed using Dada2 within QIIME2, subsequently *de novo* clustered to 97% operational taxonomic units (OTUs), and classified with a custom classifier trained on V6-V8 regions extracted from the SILVA v132 database^48–50^. Diversity and multivariate analyses were performed with Phyloseq 1.26.1^51^ and vegan v.2.5^52^. Average alpha diversities were evaluated from 100 independently rarefied datasets to the minimum read depth, using linear mixed effects (LME) models with lme4 or a two-way ANOVA^53^. Beta diversities were determined for rarefied OTU data and tested using the adonis function in vegan. Differentially abundant taxa were identified with DESeq2 v.1.22.2 and grouped by abundance pattern with DEGreport^54,55^.

#### Metatranscriptomics

Caecal content from groups of four WT or *muMT^-/-^* mice was collected at day 2 or 28 of the experiment. RNA was extracted using TRIzol (Invitrogen, Carlsbad, CA, USA) and homogenized with 0.1 mm Zirconia beads (BioSpec Products, Bartlesville, OK, USA) using a TissueLyser (QIAGEN, Hilden, Germany), followed by the PureLink RNA Mini Kit (Invitrogen, Carlsbad, CA, USA) according to the manufacturer TRIzol Plus Total Transcriptome Isolation protocol. rRNA was depleted using the QIAseq FastSelect rRNA HMR and 5S/16S/23S kits (QIAGEN, Hilden, Germany), and libraries were paired-end sequenced (150 bp) on a NovaSeq 6000 (Illumina, San Diego, CA, USA) at Genome Quebec (Montréal, Canada). Reads were processed using the MetaPro pipeline. High quality reads were mapped to the *T. mu* genome assembly using STAR v. 2.5.3a, and all remaining unmapped reads were returned to MetaPro for functional and taxonomic annotation^39,56^. Proteins with iron-related functions were predicted using FeGenie^57^. Putative cell division and cell wall biogenesis machinery was identified through sequence similarity with previously described proteins in the *E. coli* MG1655 proteome using DIAMOND v0.9.22^58,59^. DE was evaluated with DESeq2 v.1.22.2^54^. Due to read depth differences of bacterial sequences in protist-colonized and uninfected mice, reduced due to protist rRNA (Table S4), bacterial gene counts were rarefied to 600,000 reads per sample, and DE was averaged from 15 independent rarefactions. Principal component analyses were performed on DESeq2-variance stabilized counts. Functional enrichment was evaluated using EC or GO term overrepresentation as described above.

### RNA-FISH Flow Cytometry

Four million purified protists were stained with Zombie Aqua viability dye (BioLegend, San Diego, CA, USA) in PBS in the dark (1:1000, 15min, 4°C), washed twice with PBS, and the remaining steps were performed as described in the Stellaris RNA-FISH protocol for cells in suspension. After hybridization, cells were sorted on a BD LSR Fortessa X-20 cell analyzer (BD Biosciences, Franklin Lakes, NJ, USA). To test for pseudocyst formation, protists were isolated from the mouse caecum or colon and stained with wheat germ agglutinin (WGA)-FITC (Sigma Aldrich, St. Louis, MO, USA) in FACS buffer (PBS, 2% FBS, 5 mM EDTA) in the dark, either immediately or 1-3 days after anaerobic in vitro culturing. Cytospins were visualized using a Zeiss AXIO Observer microscope (Carl Zeiss AG, Jena, Germany).

### RNAScope

RNAScope was performed as per the RNAScope Multiplex Fluorescent Reagent Kit v2 (Advanced Cell Diagnostics, Newark, CA, USA) protocol. Approximately 0.5 cm cecum sections were excised from *T. mu*- colonized mice, placed in 10% neutral buffered formalin and fixed overnight at room temperature (RT) with gentle agitation. The following day, samples were washed with PBS, placed in 70% ethanol, embedded in paraffin and sliced to 7 μm sections at the Toronto Centre for Phenogenomics. Paraffin sections were baked at 40°C for 30 min in a HybEZ Oven (Advanced Cell Diagnostics, Newark, CA, USA), and treated with hydrogen peroxide at RT for 10min. Antigen target retrieval was conducted at 99°C under the 15 min standard procedure. A barrier was created around sections using an ImmEdge pen (Vector Laboratories, Burlingame, CA, USA) and allowed to dry for 15 min. Samples were treated with protease at 40°C for 30 min, and stored in saline sodium citrate solution overnight (175.3 g NaCl, 88.2 g sodium citrate, 800 mL ddH_2_O, pH 7). TSA Plus Fluorophores Fluorescein and Cyanine 3 were hybridized against protist probes TMU_00005724 and TMU_00016742 respectively, and samples were visualized using a Zeiss AXIO Observer microscope (Carl Zeiss AG, Jena, Germany).

### Transmission Electron Microscopy

Protist pellets were prepared using standard methods for the Embed 812 resin kit (Electron Microscopy Sciences, EMS)^60^. Briefly, samples were fixed with 4% paraformaldehyde, 1% glutaraldehyde in phosphate buffer (PB, 0.1M, pH 7.2), followed by 1% OsO4 in PB in the dark. After washing with PB, they were dehydrated in a 30%-100% gradient ethanol series, and infiltrated with increasing amounts of Embed 812 resin (EMS) in propylene oxide and cured in molds at 65°C. 80 nm sections were prepared with a Reichert Ultracut E microtome (Leica) on 300 mesh copper grids (EMS), and counter stained with saturated 5% uranyl acetate (EMS), followed by Reynold’s lead citrate (EMS). Sections were imaged using a Talos L120C transmission electron microscope (ThermoFisher Scientific, Waltham, Massachusetts). See supplementary methods for details.

### qPCR

qPCR of fecal DNA was performed using PowerTrack SYBR Green Master Mix (Applied Biosystems, Waltham, MA, USA). In brief, 10 ng DNA was mixed with 2X PowerTrack Master Mix and 400 nM primers, and treated as follows: enzyme inactivation at 95°C for 2 min, 40 cycles of denaturation at 95°C for 15s and annealing/extension at 60°C for 60s. Relative abundances were normalized to the universal 16S gene, and calculated using the delta-delta Ct method. Primers are provided in Table S13.

## Results

### *T. mu* colonization drives diversification of the intestinal microbiome

Colonization with *T. mu* induces profound changes in the mouse gastrointestinal tract, including the immune landscape known to maintain gut microbial homeostasis^6,19^. In light of this, we characterized the impact of *T. mu* engraftment on the local bacterial community in a healthy and immune-impaired host (Fig. 1a). We tracked bacterial composition and activities in WT and B cell-deficient (*muMt*^-/-^) C57Bl/6 mice through 16S rRNA surveys of mouse fecal pellets and metatranscriptomics of caecal microbiota (Fig. 1b, Tables S1-S3) during the first 28 days of *T. mu* colonization. Protist expansion was accelerated in *muMt*^-/-^ mice, reaching its maximum level within the first 7 days, while in WT mice, expansion steadily increased until day 28 (Fig. 1c).

*T. mu* dramatically altered gut bacterial composition in both WT and *muMt*^-/-^ mice (Fig. 1d). Bacterial richness increased in the first week of colonization, with a delay in *muMt*^-/-^ mice, before receding to near day 0 levels, while increased evenness persisted to day 28, reflecting expansions in multiple taxa (Fig. 1e). Communities in WT and *muMt*^-/-^ mice differed prior to infection, but converged toward similar profiles over the course of the experiment (Fig. 1f). Those in day 28 control mice remained similar to day 0 naïve bacterial compositions, underlining a protist-driven shift. We observed six distinct patterns of growth dynamics in *T. mu*-modulated bacteria (Fig. 1g, Table S4). The majority changed congruently in WT and *muMt*^-/-^ mice, and included taxa that increased throughout (e.g. *Bifidobacterium* and *Parabacteroides*), peaked at day 21 (e.g. *Helicobacter* and *Lachnospiraceae*), or decreased after *T. mu* colonization; five taxa exhibited discordant patterns (e.g. *Parasutterella* and *Bacteroides*). Changes in bacterial abundances were validated through qPCR (Fig. 1h).

### *T. mu* drives changes in bacterial metabolism

To investigate the impact of *T. mu* on community functions, we performed bulk RNA sequencing of mouse caecal contents at days 2 and 28 (early and chronic stages of infection) (Supplementary Fig. 1 and Table S3). 835 million reads were identified as bacterial (741 million in naïve and 94 million in colonized mice) and 157 million as *T. mu*. Colonization with *T. mu* associated with blooms in activities of Proteobacteria and Bacteroidetes (Fig. 2a). The increase in proteobacterial gene expression was accelerated in *muMT^-/-^*mice, accounting for 14% of reads by day 2 compared to 4% in colonized WT mice and 1-3% in naïve mice, suggesting a role for B cells in regulating protist-induced bacterial gene expression. The majority of these reads mapped to *Helicobacteraceae* (esp. *Helicobacter ganmani* and *Helicobacter rodentium*) previously associated with enteric inflammation and exacerbated colitis^6,15,61^.

**Figure 2.**
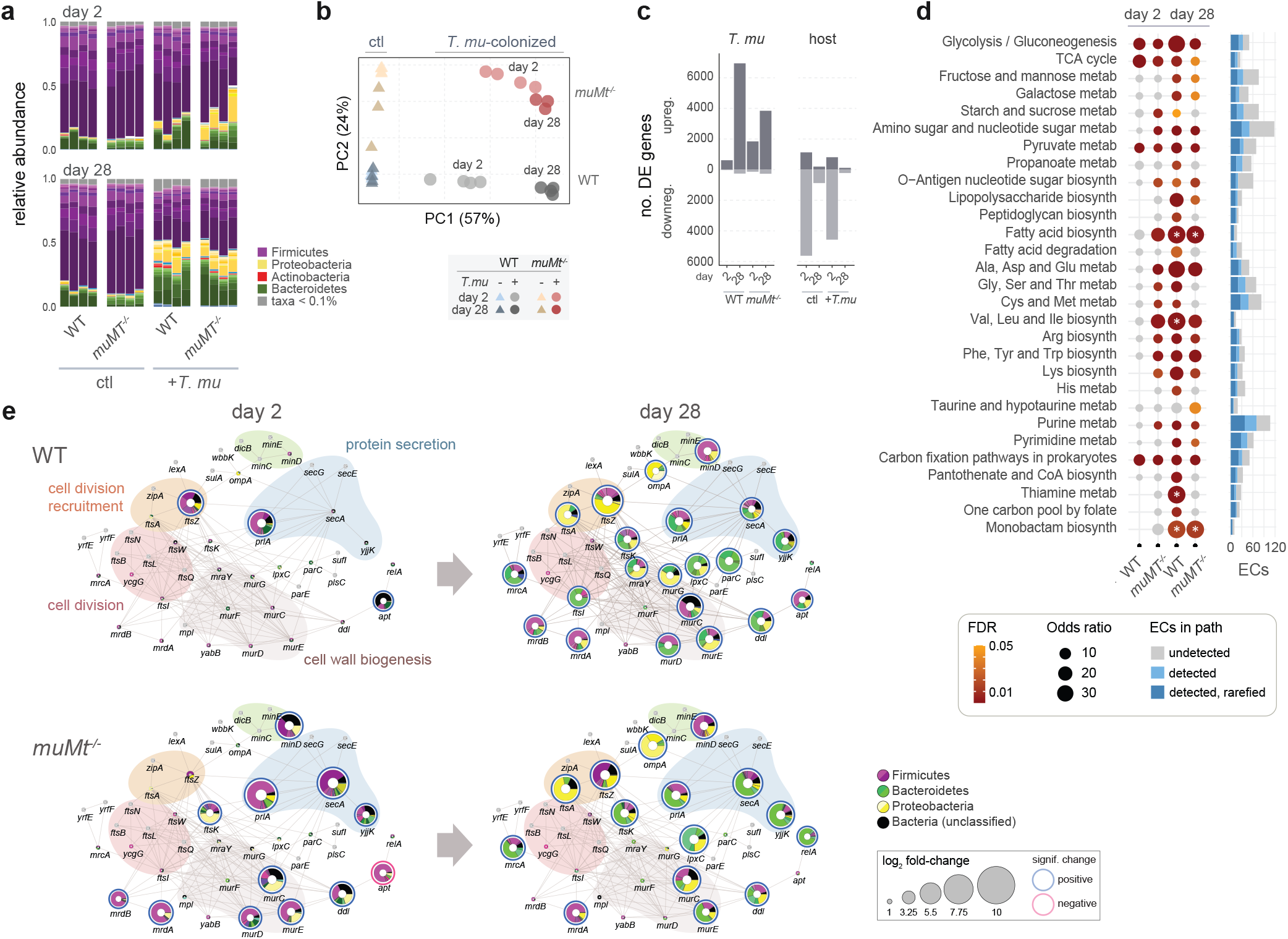
Bacterial gene expression is altered in the presence of the protist. **a**, Taxonomic profiles of putative bacterial mRNA reads. **b**, Principal component analysis of bacterial gene expression. Each circle denotes a sample isolated from a single mouse, coloured by host B cell status, presence of *T. mu* and timepoint, as shown. **c**, Numbers of significantly upregulated or downregulated bacterial genes between protist-colonized and naïve mice, or due to host B cell status, grouped as indicated (n = 3 or 4). **d**, Bacterial metabolic pathway enrichment associated with *T. mu* colonization. Enrichment was determined by the overrepresentation of EC terms in differentially expressed genes of KEGG-defined pathways. Total ECs detected in the experiment and in the rarefied gene expression matrix used in the analysis are indicated in the bar charts to the right. Significantly enriched pathways are represented as coloured dots: grey indicates *ns*, and the gradient from yellow to red represents decreasing *p* values, beginning from *p*<0.05. Sizes of dots represent odds ratios. Significance was calculated using Fisher’s exact tests. Asterisks indicate infinite odds ratios. **e**, Network of cell division, cell wall biogenesis and secretory genes, based on a previously determined *E. coli* protein network^58^. Coloured nodes represent protein homologues identified in the metatranscriptomic data, and edges are previously described protein-protein or functional interactions. Nodes in grey represent undetected genes. Genes significantly up- and downregulated are denoted with blue and red borders, respectively. Node sizes correlate with log2-fold changes between *T. mu*-colonized and naïve mice. Colours within node pie charts represent proportions of gene expression assigned to various taxa, as shown, and colours encircling groups of proteins depict functional modules as previously described.

Principal component analysis of bacterial gene expression re-capitulated the 16S data, with an overall upregulation of bacterial genes in *T. mu*-colonized mice (Fig. 2b,c). Although *muMt*^-/-^ mice exhibited higher protist-dependent upregulation at day 2, B cell-associated differences converged over time (Fig. 2c). Pathway analysis revealed upregulation of multiple metabolic pathways including glycolysis, metabolism of amino acids, and biosynthesis of peptidoglycans, polysaccharides, O-Antigen sugars and fatty acids (Fig. 2d). Metabolic activity shifted from Firmicutes at day 2 to Bacteroidetes and Proteobacteria by day 28 (Supplementary Fig. 2, 3). The expansion of Proteobacteria, particularly in the expression of succinyl-CoA synthetase (SucC) which is typically suppressed under anaerobic conditions, may indicate increased exposure to oxygen characteristic of gut inflammation and dysbiosis (Supplementary Fig.2)^62^.

Given the central role of these pathways in bacterial growth, we examined the expression of homologues to 44 genes associated with bacterial cell division and cell wall biosynthetic machinery in *E. coli* (Fig. 2e)^58^. Patterns were consistent with those described—genes were upregulated during *T. mu* colonization, with earlier responses in *muMt*^-/-^ mice and a shift from predominantly Firmicutes at day 2 to Bacteroidetes and Proteobacteria (primarily *Helicobacter*) at day 28. Day 2 bacteria from *muMt*^-/-^ mice were associated with increased chromosome segregation (e.g. *ftsK* and *ftsZ*), cell elongation (e.g. *mrdA* and *mrdB*) and peptidoglycan biogenesis (e.g. *murCDE*). By day 28, these genes were upregulated in bacteria across both hosts, together with genes involved in protein secretion (e.g. *secA*, *yjjK*, *prlA*) and porin *ompA*, which modulates infection and host immunity in Gram-negative bacteria. Collectively, these data demonstrate substantial changes in bacterial activities in response to a new protist, and a shift to an ecological niche favouring the growth of Gram-negative taxa.

### *T. mu* activity changes in a B cell-dependent manner

To monitor *T. mu* gene expression during colonization, we generated a draft assembly of the protist genome from one million PacBio sequences (median length 11,924 bp) and 8.6 million Illumina MiSeq reads. Assembly statistics are available in Tables S4 and S5. Functional annotation predicts 26,723 genes, with 999 unique GO terms, 417 enzymes and 1,932 Pfam domains (Table S6). We also identified a second distinct rRNA locus with 97% sequence identity to *T. mu*, suggesting the presence of an additional organism at lower abundance (19% of rRNA reads) (Fig. 3a). Phylogenetic comparison confirmed this novel species also belonged to the Tritrichomonadidae order (Fig. 3b).

**Figure 3.**
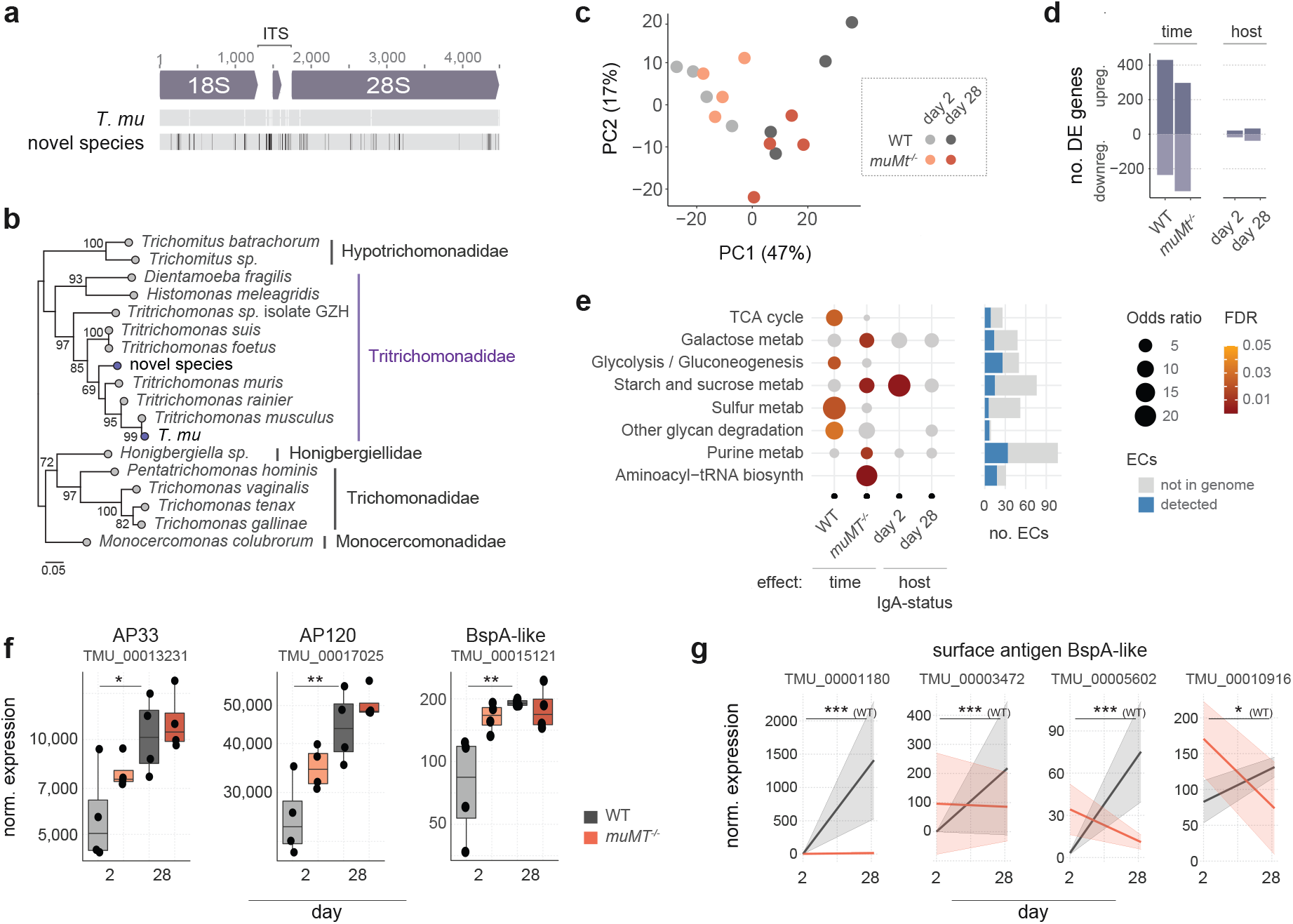
Protist gene expression changes during colonization of WT and *muMt*^-/-^ mice. **a**, Graphical depiction of aligned rRNA sequences identified in the protist metagenome assembly. The dominant sequence was synonymous with *T. mu* and mapped to 81% of reads; the second distinct sequence mapped to 19% of reads. Sequences were aligned using MUSCLE. Mismatches in the sequence of the novel organism to *T. mu* are indicated in black. **b**, Phylogeny of the two ribosomal ITS sequences identified in the metagenome. An unrooted tree was constructed from aligned sequences using the Maximum-likelihood method. Scale = number of SNPs per site. Bootstrap values >65% are shown. **c**, Principal component analysis of protist gene expression patterns at days 2 and 28, in WT or *muMt*^-/-^ mouse caeca. **d**, Numbers of differentially expressed genes associated with colonization time or host B cell status. **e**, Changes in protist metabolism during colonization. The dotplot represents enrichment of *T. mu* EC terms in KEGG-defined pathways, within sets of genes differentially expressed over time or based on host B cell status. Coloured dots represent significantly enriched pathways: the gradient from yellow to red marks decreasing *p* values, beginning from *p* < 0.05. Grey dots are *ns*. Significance was evaluated using Fisher’s exact tests, and dot sizes correlate with odds ratios. Bar charts to the right show total ECs detected in the experiment. **f**, Normalized expression of BspA-like genes showing interaction between time and host B cell status, and **g**, differentially expressed BspA-like gene and adhesins homologous to *T. vaginalis* virulence factors. Differences were evaluated using the Wilcoxon rank-sum test, and adjusted for multiple testing using the Benjamini-Hochberg approach. \**p* < 0.05, ** < 0.01, *** < 0.001, *ns*, non-significant.

We mapped 157 million metatranscriptomic reads to the protist, of which 20% mapped to coding regions (Table S3). *T. mu* gene expression changed during colonization in both WT and *muMt*^-/-^ hosts (666 and 627 DE genes, respectively), and a host B cell-modulating effect was apparent at day 28 (Fig. 3c,d). Consistent with protist growth, *T. mu* upregulated the expression of enzymes involved in energy production (e.g. the tricarboxylic acid cycle (TCA) and glycolysis) and carbohydrate metabolism (e.g. galactose, and starch and sucrose) (Fig. 3e). We probed genes with potential roles in establishing colonization: adhesins, BspA-type Leucine rich repeat regions, lectins and cysteine proteases (Tables S6-S8), protein families previously implicated in facilitating host interactions and parasite infection^31,63–66^. Homologues to adhesins AP33 and AP120, and TvBspA625, proteins with documented roles in *T. vaginalis* virulence and host cytoadherence^31,63^, were upregulated at day 28—already at day 2 in *muMt*^-/-^ mice (Fig. 3f). The expression of four additional BspA-like proteins with predicted extracellular immunogenic regions changed in B cell-dependent manners, upregulated only in WT mice (TMU_00001180, TMU_00003472, TMU_00005602, TMU_00010916; Fig. 3g, Table S9). The remaining genes showed varied changes during colonization and/or in the presence of B cells (Supplementary Fig. 4b), suggesting dynamic metabolic and accessory functions required as the protist adapts to the host environment. Future research is set to validate the role of these proteins.

### Single cell profiling of *T. mu* reveals distinct life cycle stages

The trichomonad life cycle features a variety of cell states (e.g. trophozoite, amoeboid, pseudocyst). To investigate shifts in *T. mu* cell states during colonization we carried out transcriptional profiling at the single cell level (Table S10). To further explore whether resident bacteria and B cells can influence these states, we characterized protists from a conventionalized and GF mouse, as GF mice are known to possess an immature B cell compartment^67^. Clustering of gene expression profiles from 6,000 protists (3,000 per host) revealed 15 distinct cell populations, assigned to three ‘superclusters’ (Fig. 4a). Clusters 1 through 4 (designated supercluster A) and clusters 10 and 11 were enriched in protists from the GF mouse, while clusters 5 through 7 and 15 were composed primarily of protists from the conventionalized mouse (Fig. 4b).

**Figure 4.**
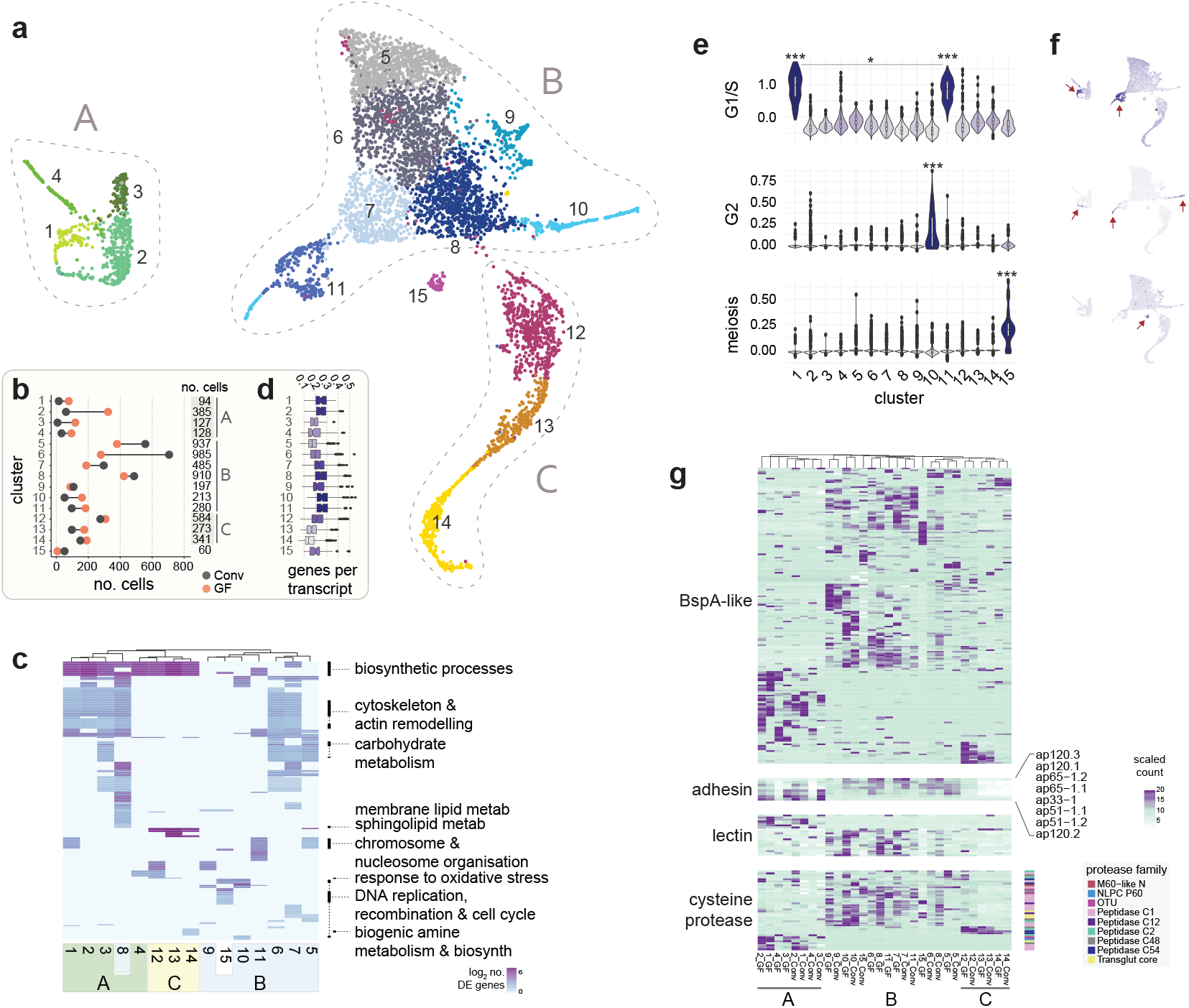
Single cell transcriptional profiling of caecal *T. mu* populations. **a**, Protist subpopulations isolated from colonized GF and conventionalized mouse caeca. Each point represents one of 6000 protists cells, clustered by gene expression profiles using UMAP dimensionality reduction, and coloured by cluster designation (3000 cells from each of two hosts). Groups of clusters, designated ‘superclusters’ A, B or C, are indicated. **b**, Numbers of cells assigned to each cluster by host are shown. Total cluster sizes are shown on the right. **c**, Heatmap of GO term enrichment within *T. mu* clusters. Rows represent significantly enriched GO terms in one or more clusters (columns). Cells are coloured and clustered by (log-transformed) numbers of upregulated transcripts per GO term. **d**, Transcriptional complexity of protist clusters, defined as numbers of genes per transcript. Boxplots indicate medians and interquartile ranges. **e**, Violin plots and **f**, feature plots depicting G1/S, G2 and meiosis scores in cells across each *T. mu* cluster. Scores were determined based on the average expression of pre-defined gene sets relative to randomly chosen control genes. Significance was evaluated using the two-sided Wilcoxon rank-sum test, \**p*<0.05, **<0.01, ***<0.001. Asterisks above clusters indicate significantly higher score relative to all other clusters. Lines denote specific comparisons between two clusters. **g**, Heatmap showing expression of (*top* to *bottom*) BspA-like proteins, putative adhesin proteins, lectins and Cysteine proteases across protist clusters. Colours to the right of Cysteine proteases indicate the protein family (Pfam).

Functional enrichment of GO terms suggested the presence of actively metabolizing cells, protists at various stages of cell cycle, and those undergoing pseudocyst formation (Fig. 4c). Clusters 1-3 and 5-8, for example, were enriched in cytoskeleton & actin remodeling, carbohydrate metabolism and biosynthetic processes and may represent actively feeding and growing *T. mu* trophozoites. Cluster 8 is enriched in sphingolipid metabolism, which contributes to regulation of *Giardia* encystation^68^. Since cluster 14 is enriched in biogenic amine metabolism, a pathway associated with encystation in *Entamoeba invadens*^69^, and exhibits lowest transcriptional complexity (Fig. 4d), we suggest these cells represents a pseudocyst form of *T. mu*^70,71^. Furthermore, the upregulation of oxidative stress response in the neighbouring clusters 12 and 13 suggests the encystation transcriptional program may be triggered through stress-induced pathways.

Enrichment of chromosome reorganization, DNA replication and cell cycle checkpoint in clusters 1, 5, 10 and 11 identified protists undergoing cell cycle. While cluster 15 shared cell cycle terms with these clusters, it was uniquely enriched in DNA recombination, indicative of sexual or parasexual replication. To investigate further, we generated functional enrichment scores for the expression of *T. mu* homologs associated with the G1/S and G2 stages of the cell cycle, and those associated with meiosis (Tables S6 and S7)^30,72^. Scores for expression of G1/S marker genes were highest in clusters 1 and 11, and cluster 10 scored highest for the G2 phase (Fig. 4e,f; Supplementary Fig. 5a,b). Cluster 15 scored highest for meiosis-related markers (Fig. 4f). In addition to meiosis-specific genes *dmc1*, *hop2A* and *mnd1*, required genes *rad1*, *mre11*, *smc2*, *smc3* and *smc5* were also significantly upregulated in cluster 15 (Supplementary Fig. 5c), consistent with sexual or parasexual replication. Of the 60 cells associated with this this cluster, only 8 were from the GF mouse. Together with other differences in cluster distributions between conventionalized and GF mice (Fig. 4b), our results suggest that the constituent microbiome may play a role in modulating the protist life cycle. However, we note that despite purification and antibiotic treatment of the protist, we detected bacterial DNA in the GF mouse, limiting our conclusions on microbiome impact.

### Protist populations express distinct sets of virulence genes

Since the distribution of protist cell states was altered in the GF mouse, we investigated how these changes might impact the expression of genes implicated in colonization (BspA-like proteins, adhesins, lectins and cysteine proteases; Fig. 4g). Many genes exhibited cluster and/or host-specificity, with profiles broadly defined by the three major superclusters (Table S11). Notably, cells in cluster 15 (the putative sexual stage) expressed distinct BspA-like genes in GF and conventionalized hosts, implicating a role for the microbiome in regulating the protist life cycle, while putative encysting stages (clusters 12-14) shared similar expression patterns in both mice. Of the nine putative adhesins, which mediate binding to host epithelia in *T. vaginalis*^31^, we detected expression for eight (Fig. 4g). These genes were absent from pseudocyst-associated clusters 13 and 14, supporting their role in colonization. Lectins were depleted in clusters 12-14, as well as the putative sexual stage, cluster 15. Of the four cysteine proteases expressed in putative pseudocysts, two are members of the C54 peptidase family which have previously been implicated in cell starvation and differentiation^73,74^. These data present a dynamic arsenal of virulence factors, exhibiting cell stage and host-specific expression, potentially modulated by resident microbiota, which may mediate protist colonization and transmission.

### Distinct *T. mu* cell stages are detected *in vivo*

We validated the presence of distinct *T. mu* populations in the mouse intestine through *in situ* fluorescent labelling of cluster-specific mRNAs. Microscopic imaging of caecal sections from WT mice revealed protists as large, nucleated cells restricted to the gut lumen (Fig. 5a). Protists expressing one or more cell-stage specific gene could be visibly identified in the cecum (e.g. TMU_00005724, specific to cells of supercluster B, and/or TMU_00016742 expressed in a subset of B; Fig. 5b,c). Fluorescent *in situ* hybridization flow cytometry (FiSH-Flow) revealed expression of TMU_00005724 in all caecal protists (Fig. 5d). Consistent with cluster-specific profiles, expression of TMU_00016742 (clusters 5, 6 and 9) exhibited higher overlap with TMU_00005724 than did TMU_00009244 (cluster 12) and TMU_00001185 (cluster 14) (Fig. 5c,e). *muMT^-/-^* mice harbored more TMU_00016742-expressing protists, suggesting B cell influence on the *T. mu* transcriptional program (Fig. 5e). Since TMU_00001185 was unique to cells of cluster 14 (predicted pseudocysts) and pseudocyst formation occurs during host egress, we tested for its expression in protists freshly isolated from the caecum, colon, and for up to 3 days of *in vitro* culturing (Fig. 5f). As expected, caecal isolates contained fewest TMU_00001185-expressing cells, and their proportions increased during *in vitro* culturing. Supporting their identity as pseudocysts, the same pattern was observed for presence of chitin, a critical component of cysts and pseudocysts, stained using fluorophore-labelled wheat-germ agglutinin (WGA) (Fig. 5g, Supplementary Fig. 6)^71,75,76^. WGA-labelled protists furthermore showed thicker cell walls, consistent with cyst-like structures (Fig. 5h). The *in situ* data confirm the presence of transcriptionally distinct groups of protists in the cecum, and illustrate the utility of cluster-specific genes to monitor *T. mu* life cycle dynamics during colonization.

**Figure 5.**
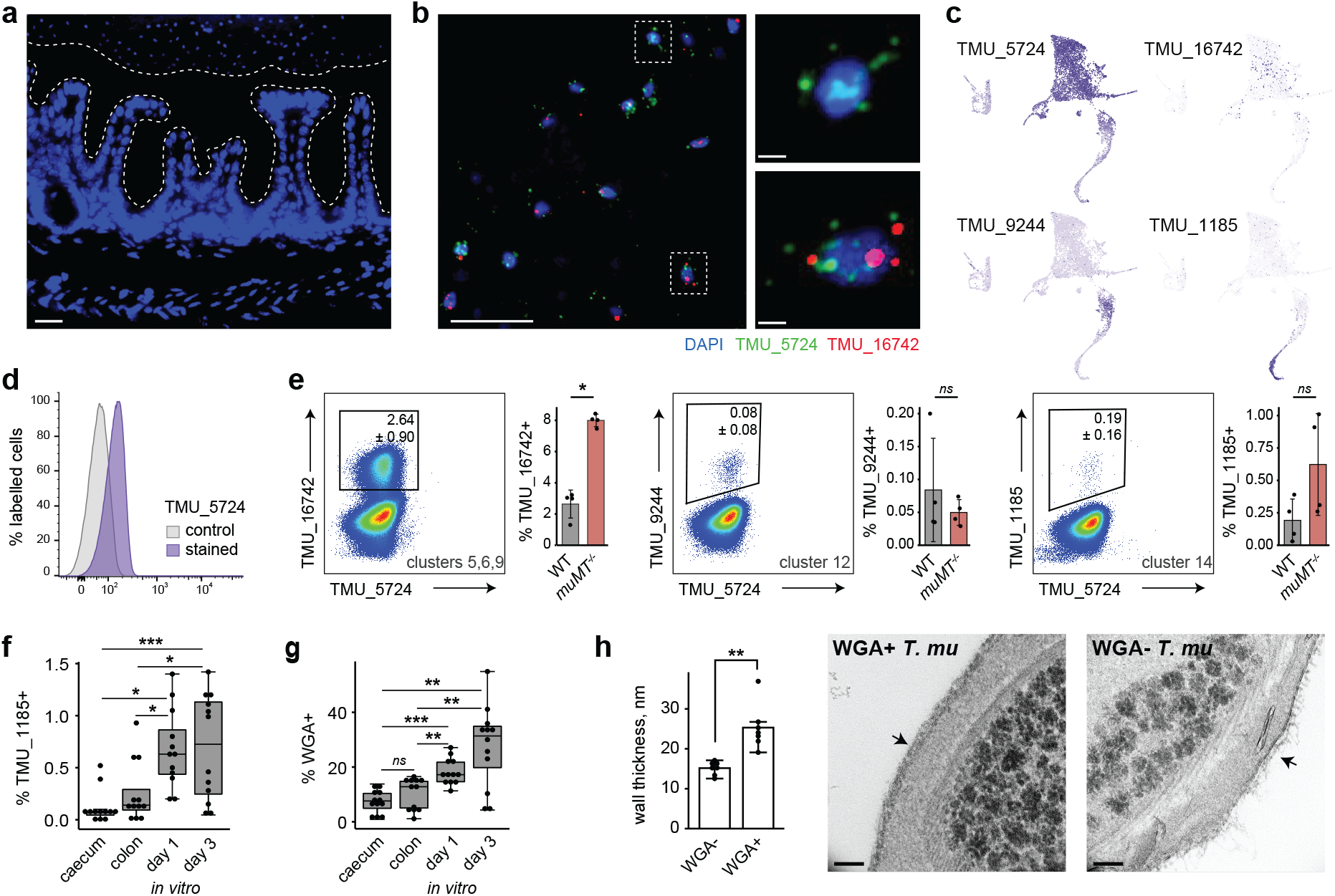
*T. mu* subpopulations are detected *in situ*. **a,** Immunofluorescence micrograph of mouse cecal tissue after colonization with *T. mu* for 21 days. Nuclear staining (blue) shows host and protozoan DNA, and white dotted lines denote the mucus layer. Scale bar = 10μm. **b**, RNAscope immunofluorescence of *T. mu* transcripts TMU_00005724 (green) and TMU_00016742 (red) on sections of cecal tissue. Nuclear DNA is stained blue. Left image scale bar = 10μm; right images scale bars = 1μm. **c**, Feature plots showing the expression of cluster-specific genes. **d**, FISH-Flow analysis of freshly isolated *T. mu* at 21 days post colonization. Histogram shows control unstained cells (grey) and cells stained with fluorescent probes for TMU_00005724 (purple) **e**, FISH-Flow analysis of *T. mu* isolated 28 days post colonization from WT or *muMt^-/-^* mice. Numbers adjacent to gates represent the average percentage +/-standard deviation of probe-expressing *T. mu*. Data shown is representative of 4 independent experiments, and adjacent plots show results from all experiments. Significance was tested using the Wilcoxon rank-sum test. **f**, Percentages of TMU_00001185 probe-positive or **g**, WGA-FITC stained protists freshly isolated from WT mouse caeca and colons, or cultured *in vitro* for one and three days. n=3 animals or culture plates per group, from four independent experiments. Significance was tested using one-way ANOVA, and adjusted for multiple comparisons using Tukey’s test. **h**, Cell wall thickness of FACS-sorted protists hybridized or not to WGA-FITC. Cells were imaged using TEM, and cell walls were measured for all cells (5-10) across 7 view fields per group. Scale bars, 100 nm. Bars are means and whiskers show interquartile ranges. \**p* < 0.05, \*\**p* < 0.01, \*\*\**p* < 0.001, *ns* not significant.

### *T. mu* and gut bacteria compete for nutrients

As *T. mu* colonizes the gut, it must compete with resident microbiota and the host for nutrients. Since trichomonads require high concentrations of iron for growth^77,78^, we hypothesized that its consumption by *T. mu* would exert pressure on bacterial iron acquisition and storage systems. Metatranscriptomic data confirmed increasing protist iron consumption through the upregulation of ferredoxin-1 (TMU_00007489), an enzyme required for energy production, and also adhesin TMU_00017025 with predicted pyruvate:ferredoxin oxidoreductase activity (Fig. 6a,3f)^79,80^. Both enzymes were widely expressed across protist cell states, but reduced in clusters 13 and 14, consistent with lower energy requirements of a pseudocyst (Fig. 6b). The upregulation instead of ferredoxins TMU_00018447 and TMU_00005891 in clusters 12 and 14 may be specific to encysting forms, and potentially involved in mediating stress response. Similar expression of these genes in conventionalized and GF mice suggests *T. mu* is agnostic in terms of iron acquisition to the resident microbiome. In response to *T. mu*, expression of bacterial iron acquisition and storage systems increased, including siderophore synthesis and transport genes (Fig. 6c,d), predominantly in *Helicobacter*, *Bacteroides*, *Parabacteroides* and *Mucispirillum schaedleri* (Supplementary Fig. 7b). Host B cell status minimally impacted these systems, suggesting more direct competition between *T. mu* and resident microbiota (Supplementary Fig. 7c).

**Figure 6.**
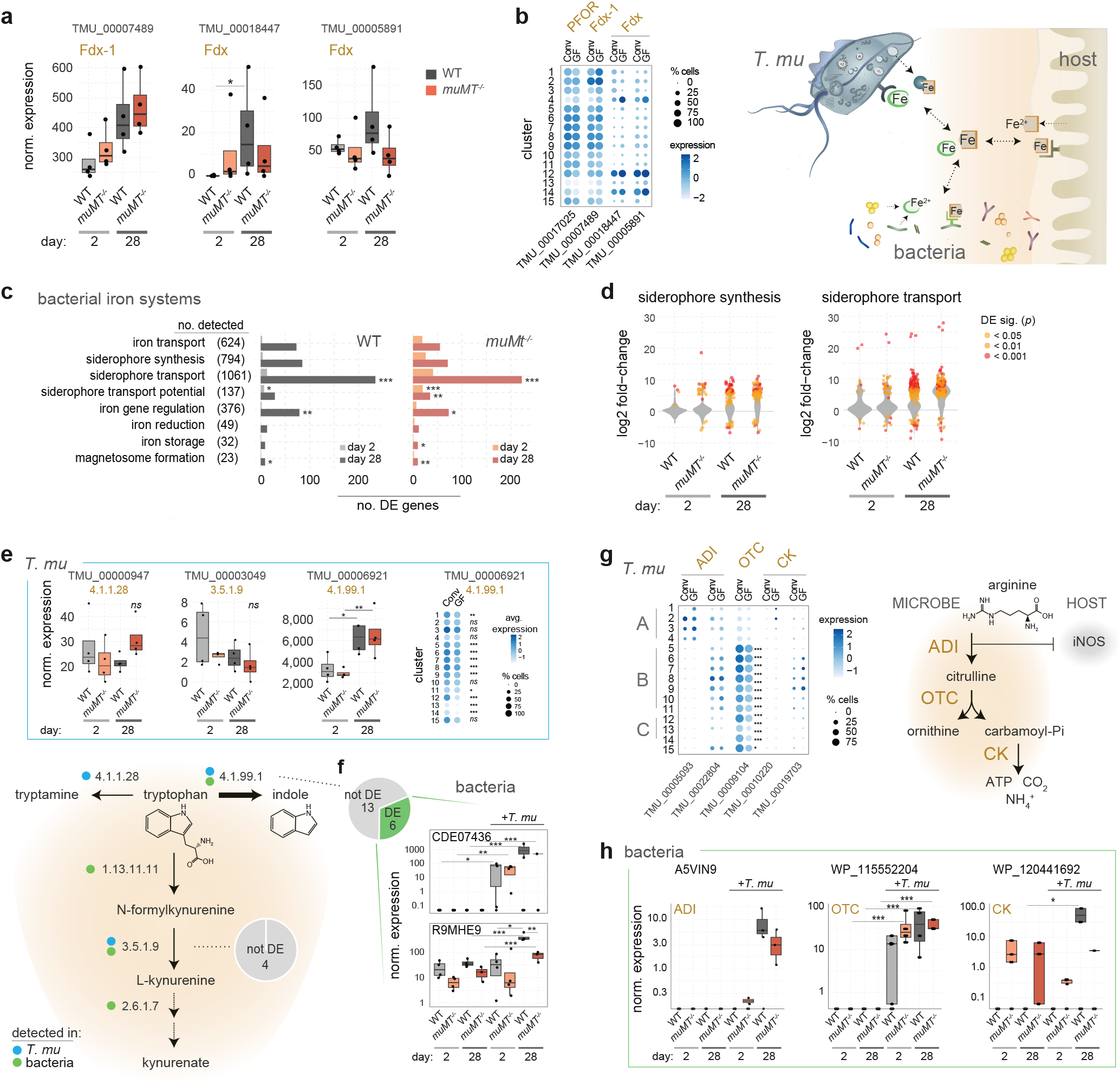
*T. mu* and resident bacteria compete for resources in the gut lumen. Shown are changes in expression of bacterial and *T. mu* genes associated with iron, **a**-**d**, generation of tryptophan catabolites, **e** and **f**, and the arginine dihydrolase pathway, **g** and **h**. **a**, Expression of putative *T. mu* ferredoxin (Fdx) genes in metatranscriptomic data at days 2 and 28. **b**, PFOR and Fdx gene expression among protist subpopulations isolated from GF and conventionalized mice. Colour intensity correlates with average expression across cells and dot sizes represent percentages of cells in each cluster with detected expression. **c**, Enrichment of iron-related gene expression in bacteria, in the context of *T. mu* colonization. Shown are numbers of DE genes per iron-related gene family in *T. mu*-colonized versus naïve mice at days 2 and 28. Total numbers of genes identified in each family in the metatranscriptomic data are indicated in brackets. Significance of enrichment was tested using the hypergeometric test. **d**, Median log2 fold-change of bacterial siderophore transport and synthesis gene expression between colonized and naïve mice. Significantly DE genes are shown as coloured points: the gradient from yellow to red marks decreasing *p* values, beginning from *p*<0.05. **e**, Expression of *T. mu* and **f**, the most highly expressed bacterial (read count >100) enzymes with predicted function in the production of tryptophan metabolites. Dotplots in E show gene expression among protist subpopulations, as above. Pie charts in F represent the total numbers of tested and DE bacterial enzymes. Blue and green dots in the graphic show enzyme activities identified in *T. mu* and/or bacterial genes, respectively. **g**, Expression of *T. mu* enzymes associated with the arginine dihydrolase pathway among protist subpopulations isolated from GF conventionalized mice, as above. **h**, Expression of bacterial enzymes associated with the arginine dihydrolase pathway. Shown are the only ADI present in the rarefied gene count matrix, and the two differentially expressed OTC and CK genes. Gene expression in metatranscriptomics data was evaluated using DESeq2, and in scRNA-Seq data using Seurat. All expression values are derived from normalized gene counts. Boxplots and violin plots show medians and interquartile ranges. *P* values were corrected for multiple testing using the Benjamini-Hochberg approach. \**p*<0.05, **<0.01, ***<0.001

We also tracked expression of genes associated with tryptophan and arginine metabolism, amino acids implicated in host:microbiome interactions^81,82^. The protist tryptophanase TMU_0006921 (EC 4.1.99.1) was upregulated at day 28 (Fig. 6e). As enzymes in competing pathways (conversion of tryptophan to kynurenate and tryptamine) exhibited minimal expression with no temporal shift, *T. mu* appears to preferentially drive production of indole (Fig. 6e). Remarkably, the two most highly expressed bacterial tryptophanases were also upregulated in *T. mu* colonized-mice suggesting bacteria further promote the indole production in the presence of the protist (Fig. 6f). Consistent with this, scRNA-Seq data showed upregulation of *T. mu* tryptophanase in the conventionalized mouse and reduction in putative pseudocyst clusters (13 and 14) suggesting reduced dependence on host mucosal homeostasis (Fig. 6e).

Arginine is an important source of energy for both trichomonads and bacteria. Its depletion by microbes through the arginine dihydrolase pathway has an additional immunomodulatory role, limiting host production of antimicrobial NO^81,82^. None of the three *T. mu* enzymes in the associated pathway: arginine deiminase (ADI), ornithine transcarbamylase (OTC) or carbamate kinase (CK), were significantly associated with either colonization time or host B cell status, suggesting continual arginine consumption (Supplementary Fig. 4d). We did note selective expression across protist cell states (Fig. 6g). Conversely, *T. mu* colonization was associated with increased expression of several bacterial arginine dihydrolase genes, including a Lactobacillales ADI and *Helicobacter* OTC (Fig. 6h).

The dynamics of the acquisition and metabolism machinery for these key gut nutrients suggests competitive as well as cooperative relationships between the protist and the resident microbiota.

## Discussion

Previous work has revealed that protists mediate antagonistic and mutualistic interactions with intestinal bacteria with consequences for the host, primarily through studies of microbial composition^6,7,15,83,84^. Here we demonstrated interactions between *T. mu* and resident microbiota at a functional level in a healthy and immunodeficient host, and identified genes predicted to facilitate protist colonization. We showed that, consistent with a previous study, *T. mu* induces longitudinal shifts in bacterial composition^17^. However, our study notes increased bacterial diversity and importantly the abundance and activity of *Helicobacter* spp. in the presence of *T. mu*, reported also during *T. foetus* colonization^85^. The shift from Firmicutes to Bacteroidetes and Proteobacteria in colonized mice suggests environmental pressure favouring Gram negative taxa. As Gram negative bacteria harbor surface LPS capable of boosting immunity, their bloom may contribute to the intestinal immune activation observed during *T. mu* engraftment^6,86^. The earlier occurrence of these changes in B cell-deficient mice suggests a failure to control commensal bacteria that might otherwise be mediated through *T. mu*-induced IgA reactivity^19,87,88^. We speculate this reactivity might serve to limit adverse responses by the bacteria as the protist attempts to colonize. Similarly, the accelerated expansion of *T. mu* and earlier upregulation of its adhesins and TMU_00015121, a homologue of the antigenic *T. vaginalis* BspA625 protein, signals changes in protist immunogenicity and suggests that B cell surveillance plays a critical role during early protist infection^63,89^.

Within our scRNASeq data, we identified distinct protist populations representing different stages of the *T. mu* life cycle. Cluster-specific patterns of expression of protein families implicated in host cell binding or parasite virulence suggest the protist alters its host adhesive properties or antigenicity as it transitions through its life cycle^31,63,65,66^. *T. vaginalis* and *T. foetus* exist in one of three forms—actively growing trophozoites, host-adherent amoeboid cells and environmentally resistant pseudocysts^70,71,90–92^. The capture of actively metabolizing *T. mu* in the lumen, together with cells exhibiting reduced activity (with little to no expression of adhesins) and stress-activated pathways suggests our data captured the trophozoite and pseudocyst states. Supporting the ability of *T. mu* to penetrate the intestinal mucous layer and gain access to host epithelia, we identified a large repertoire of expressed cysteine proteases and enzymes of the N-glycan degradation pathway, capable of degrading O-glycans, the dominant glycans found in mucins^93,94^.

We validated the pseudocyst stage using cluster-specific and chitin markers on mouse intestinal contents and *in vitro* cultures, and TEM of purified protists. A similar transcript-labelling approach could be used to capture the amoeboid state through exposure to host epithelia^90,91^. One possibility is that we have already captured the amoeboid state in our scRNA-Seq data, in supercluster A. The majority of protists in these populations were recovered from the GF mouse, which are known to have a more penetrable intestinal mucus^95^. These cells express different suites of adhesins, cysteine proteases and enzymes involved in arginine metabolism. Their identification has clinically relevant implications in a diseased or inflamed gut where the protist may further exacerbate symptoms^6^. The discovery of meiosis-specific genes is particularly intriguing as it suggests the presence of sexual replication, with implications for host adaptation and immune evasion. Sexual recombination has previously been hypothesized in *T. vaginalis* and *G. duodenalis*^30,96–99^. While our data may suggest that the microbiome promotes meiosis (as noted by the low proportion of meiotic cells in the GF mouse), we did detect bacterial DNA in the GF mouse. Differences in meiotic cells may therefore arise from timing of microbiome exposure. Although unlike the GF mouse, the conventionalized mouse was exposed to commensal bacteria for four weeks prior to *T. mu* infection, which would have affected intestinal morphology and immune cell differentiation, additional experiments are needed to confirm the role of the microbiome on the *T. mu* life cycle^100^. To date, attempts at axenic culturing of *T. mu* have proven challenging.

To successfully colonize, *T. mu* must compete with resident bacteria for nutrients. Parasitic trichomonads are known to require high quantities of iron to sustain their growth, whereas low iron conditions hinder their adhesion to host epithelia^78,101^. To limit growth and virulence of invading pathogens, hosts restrict access to luminal iron through sequestration by proteins such as lactoferrin^77,78,102^. The upregulation of bacterial iron acquisition machinery in response to *T. mu* has the potential, therefore, to regulate protist growth and virulence, an effect which may be lost in a context of surplus iron provided for example by nutritional supplements^5^. We also noted synergistic increased expression of genes metabolizing tryptophan to indole in *T. mu* and bacteria. Production of indole by *T. foetus* and *T. vaginalis* was shown to impact host immunity and barrier function, and bacterial signalling^103–105^. Recently there has been much interest in the production of neuroactive kynurenines from tryptophan by the gut microbiome^106,107^. Through promoting production of indole at the expense of alternative products, tryptamine and kynurenine, our findings question whether protists similarly modulate the gut-brain access.

The ecological shift induced by *T. mu* identifies it as a keystone species of the murine gut microbiome. Applying similar sequence-based and *in situ* methodologies will provide insight into how other gut protists might impact gut health.

## Acknowledgements

This work was funded by the Canadian Institutes of Health Research (MRT-168043) to J.P., M.E.G. and A.M; the Natural Sciences and Engineering Research Council (RGPIN-2019-06852) to J.P.; the Intramural Research Program of the National Institute of Allergy and Infectious Diseases at the National Institutes of Health to M.E.G.; a Restracomp scholarship administered by the Research Training Centre (Hospital for Sick Children) and a graduate scholarship from the Government of Ontario to A.P. A.M. is supported by the Canadian Foundation for Innovation John R. Evans Leaders Fund, the Canadian Institutes of Health Research (PJT-388337, PJT-480765) and a Natural Sciences and Engineering Research Council (RGPIN-2019-04521). A.M. is the Tier 2 Canadian Research Chair in Mucosal Immunology and supported by the Tier 2 CRC-CIHR program (CRC-2021-00511). Computing resources were provided by the SciNet High Performance Computing (HPC) Consortium; SciNet is funded by the Canada Foundation for Innovation under the auspices of Compute Canada, the Government of Ontario, Ontario Research Fund - Research Excellence, and the University of Toronto. We thank the Temerty Faculty of Medicine Microscopy Imaging Lab at the University of Toronto for TEM sample preparation and training.

## Author contributions

J.P., A.M., M.E.G., and A.P. conceived and designed the study. A.P. and A.K. isolated nucleic material for high throughput sequencing, and A.P. performed sequence data analyses. E.Y.C. performed mouse colonization experiments, performed flow cytometry and microscopic analyses. N.N. performed protozoan enzyme annotation. N.A. and J.H. participated in sample collection and *in vitro* culturing experiments. E.C.A.F. generated the phylogenetic tree. A.P., J.P., A.M. and M.E.G. wrote the manuscript. All authors reviewed and/or edited the paper.

## Data Availability Statement

Sequence data generated in this study have been deposited to the NCBI Sequence Read Archive under the BioProject identifiers PRJNA913581 and PRJNA914770.

