## Supplementary information for "The commensal protist *Tritrichomonas musculus* exhibits a dynamic life cycle that induces extensive remodeling of the gut microbiota"

**Supplementary Methods**

**Genome sequencing and annotation**

Genomic DNA was extracted from 100 million sorted protists using the MagAttract HMW DNA Kit (QIAGEN, Hilden, Germany). Sequencing libraries were prepared and sequenced using PacBio Sequel technology on two SMRT cells at the McMaster University Farncombe Metagenomics Facility (Hamilton, Canada). Reads were error corrected using CANU v1.8^71^, assembled into contigs using Flye v.2.4.1^72^, and the contigs were subsequently polished with eight million 300 bp paired-end Illumina MiSeq reads, sequenced at the National Institute of Allergy and Infectious Diseases (Bethesda, Maryland) using BWA^73^ and Pilon v.1.23^74^. Annotation was carried out with Maker v2.31^75^ as follows. Repetitive regions were identified *de novo* and masked using RepeatModeler v.1.0.11^76^ and RepeatMasker v.4.0.7^77^. Transfer RNA genes were predicted with tRNAscan-SE v2.0.6^78^. Gene models were predicted *ab initio* using two rounds of training with SNAP v.2013-11-29^79^, followed by Augustus v.3.3.1^80^ with BUSCO v.2.0.1^81^. Reference sequences included related trichomonad *T. foetus* expressed sequence tag (EST) library (CX159216.1), *T. foetus* strain K (3AUP000179807) and *Trichomonas vaginalis* G3 (3AUP000001542) proteomes, and *Pentatrichomonas hominis* and *Dientamoeba fragilis* protein sequences available in the NCBI nr database (accessed August 21, 2019). Functional annotation was performed with InterProScan v.5.30-69.0^82^, the HmmerWeb v.2.41.2^83^ hmmscan algorithm (E-value ≤1e-05) and Architect^84^ (confidence ≥0.5). Genes encoding adhesins, meiosis and cell cycle-related proteins were identified based on sequence homology with *T. vaginalis* proteins^26,35^ retrieved from the TrichDB database^85^ using BLAST^86^ (E-value 1e-5, 30% sequence identity and 40% query coverage cut-offs). The genome assembly is available at: <https://github.com/ParkinsonLab/Tritrichomonas-murine-microbiome-interactions/draft-genome-assembly>.

**Transmission Electron Microscopy**

Protozoa pellets were prepared using the standard methods for the Embed 812 resin kit (Electron Microscopy Sciences, EMS). (Hayat, M. A. Principles & Techniques for Electron Microscopy. Second Edition (1981)). Briefly, samples were fixed with 4% paraformaldehyde, 1% glutaraldehyde in phosphate buffer (PB; 0.1M, pH 7.2) for 1 hour at RT and overnight at 4°C, washed 3x with PB, and secondary fixed with 1% OsO4 in PB in the dark. Samples were washed again 3x with PB for 10 min at RT. Samples were dehydrated in a gradient ethanol series: 30% ethanol for 15 min, 50% ethanol for 20 min, 70% ethanol for 30 minutes, 90% ethanol for 45 minutes, and 100% ethanol for 60 min. Samples were infiltrated with the Embed 812 resin kit (EMS) diluted with propylene oxide: 100% propylene oxide for 20 min, 33% (v/v) Embed 812 resin mixture in propylene oxide for 2 hours, 67% (v/v) Embed 812 resin mixture in propylene oxide for 3 hours, 100% Embed 812 resin mixture overnight, fresh 100% Embed 812 resin mixture for 2 hours. After infiltration samples in resin were put in molds and cured at 65°C for 48 hours.

Resin blocks were sectioned to 80 nm thickness with a Reichert Ultracut E microtome (Leica), collected on 300 mesh copper grids (EMS), and counter stained for 10 min each using saturated 5% uranyl acetate (EMS), followed by Reynold’s lead citrate (EMS). Prepared grids were placed on a filter paper mat in labelled Petri dishes and stored in a desiccator until imaging. The sections were imaged using a Talos L120C transmission electron microscope (Thermo Scientific) equipped with a BM-Ceta scientific CMOS camera at an accelerating voltage of 120KV.
