## Supplementary Fig for "The commensal protist *Tritrichomonas musculus* exhibits a dynamic life cycle that induces extensive remodeling of the gut microbiota"

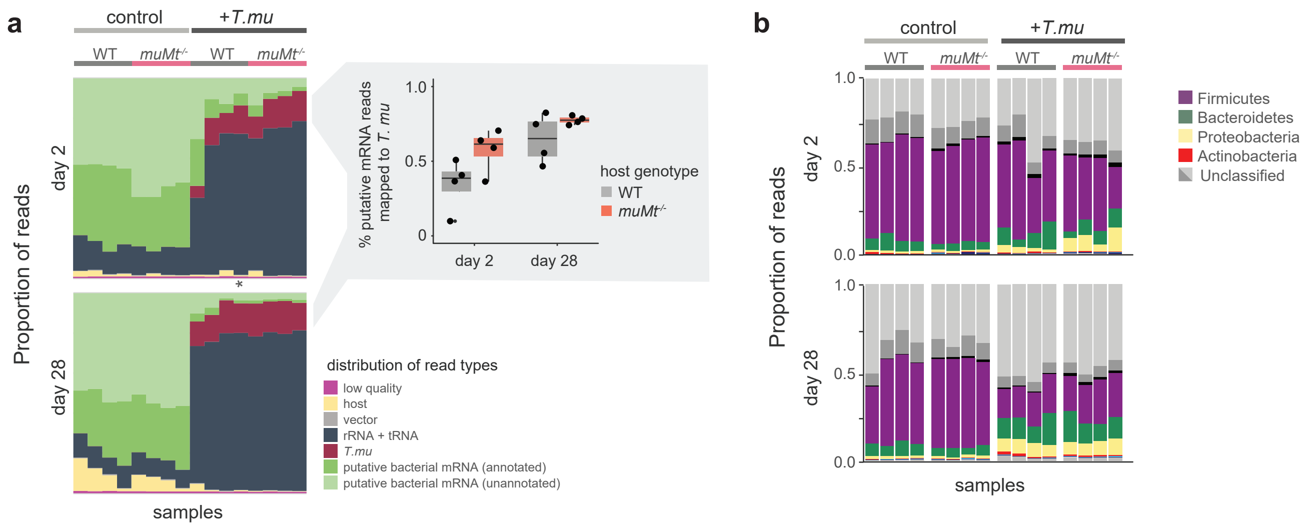


**Supplementary Figure 1.** Caecal metatranscriptomics of *T. mu*-colonized and naïve (control) WT and *muMt*$^{-/-}$ mice. **a**, Breakdown of caecal RNA reads from filtering and annotation steps. Columns represent samples from individual mice. Outset graph to the right shows percentages of putative mRNA reads mapped to the *T. mu* genome assembly. **b**, Taxonomic classification of putative bacterial transcripts.

Glycolysis/Gluconeogenesis, pentose phosphate and TCA

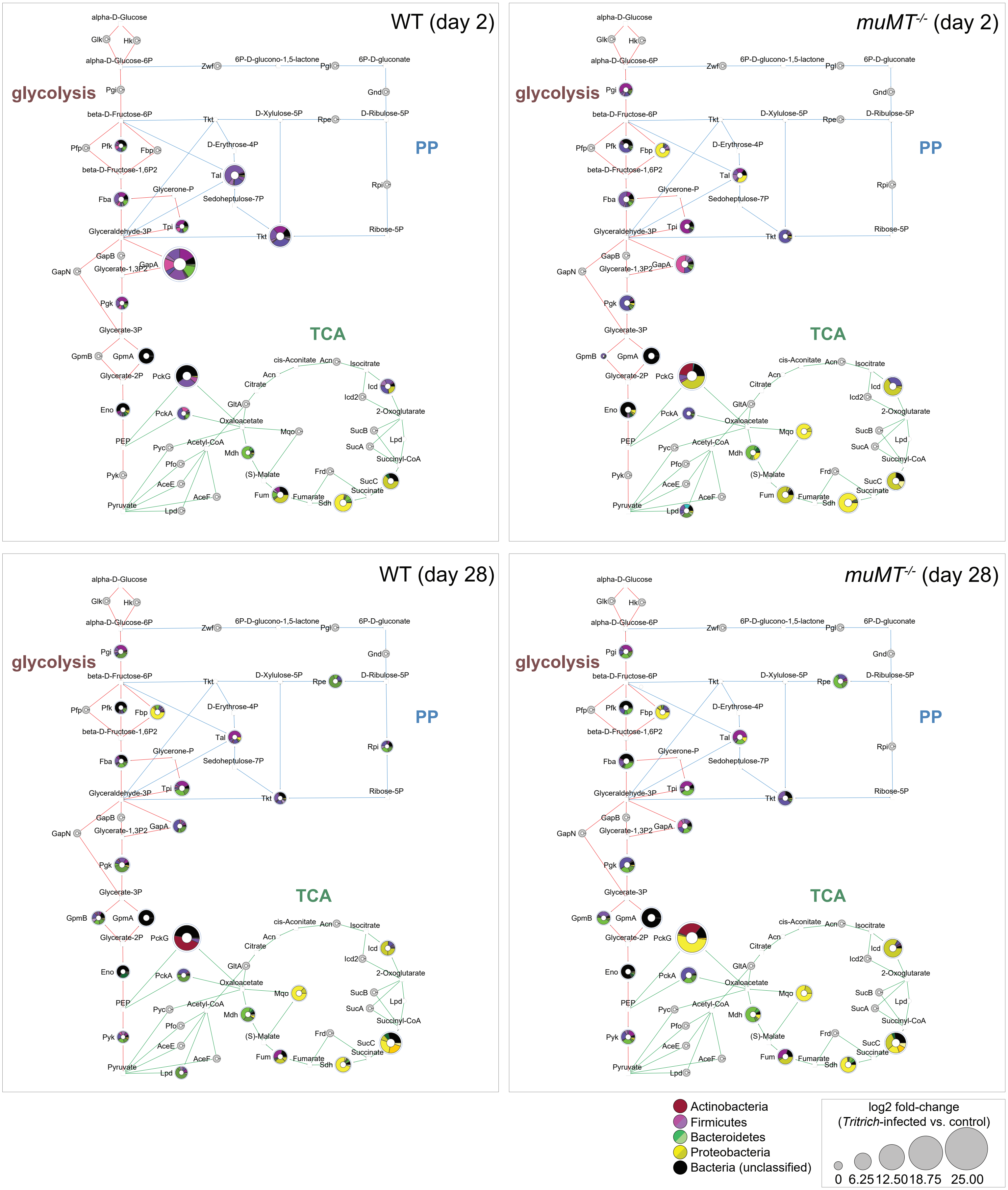


**Supplementary Figure 2**. Upregulation of bacterial metabolism in response to protist colonization. Depicted are glycolysis/gluconeogenesis (ec00010), tricarboxylic acid (TCA) cycle (ec00020) and the pentose phosphate (PP) (ec00030) pathways in gut microbiota after 2 or 28 days of infection in WT or B-cell deficient (*muMT-/-*) hosts. Genes significantly up- and downregulated ($p < 0.05$ in DESeq2 analysis) are indicated with blue and red borders, respectively. Sizes of nodes represent log2-fold changes between *T. mu*-colonized and uninfected control mice (n=4 per group). Pie charts depict the phylogenetic source of the gene expression as follows: yellows represent Proteobacteria (dominated by *Helicobacter*); shades of green are Bacteroidetes (dominated by *Bacteroides* and *Parabacteroides*); pinks and purples are Firmicutes (dominated by Lachnospiraceae and *Clostridium*); black represents unclassified bacteria.

Valine, leucine and isoleucine biosynthesis (ec00290)

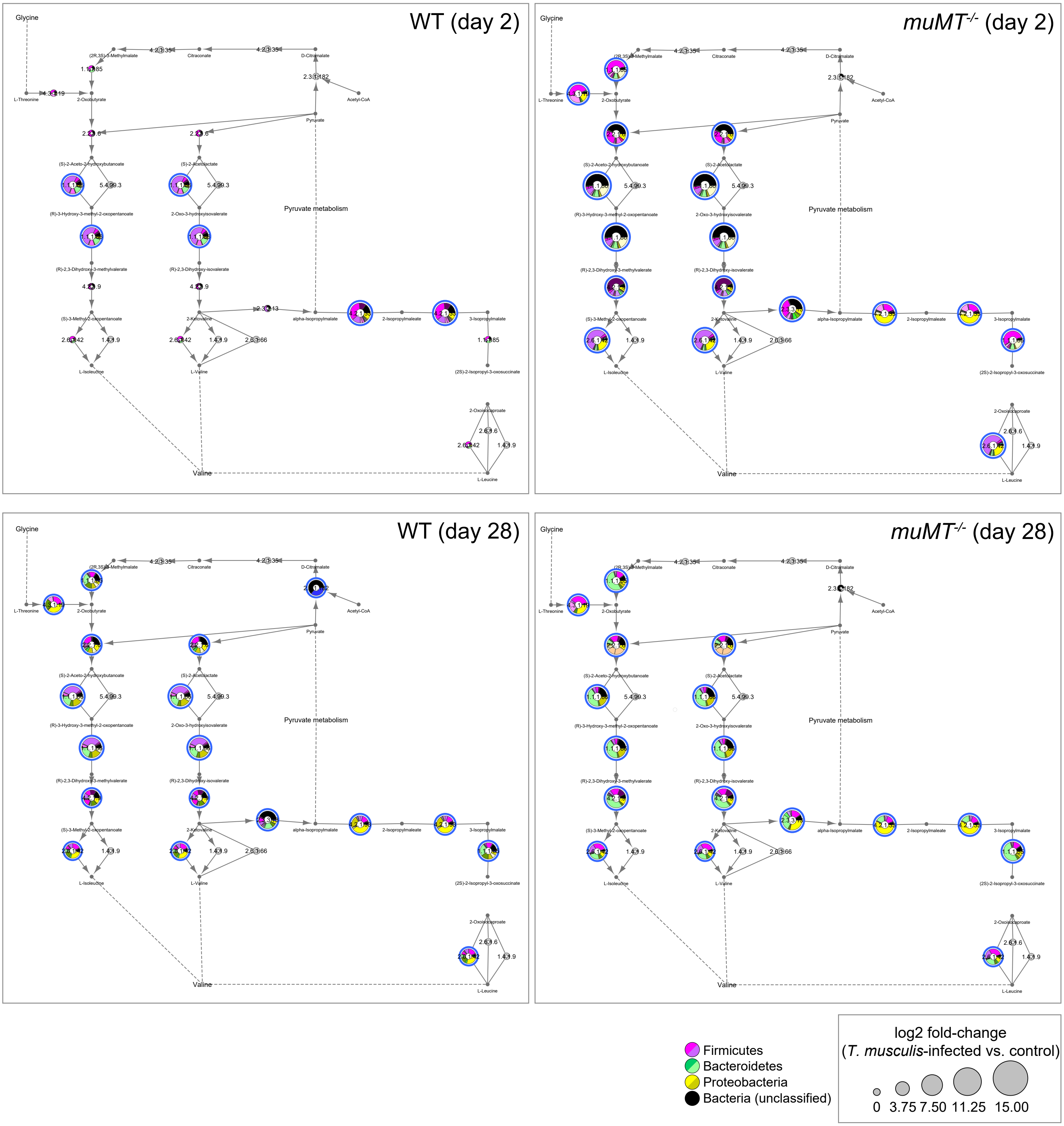


**Supplementary Figure 3**. Upregulation of Valine-Leucine-Isoleucine biosynthesis in mouse cecal microbiota in response to four weeks of protist colonization. Depicted is metabolic pathways protein interaction network based on Genes significantly up- and downregulated ($p < 0.05$ in DESeq2 analysis) are indicated with blue and red borders, respectively. Sizes of nodes represent log2-fold changes between *T.mu*-colonized and uninfected control mice at day 28 of the experiment. Pie charts depict the phylogenetic source of the gene expression as follows: yellows represent Proteobacteria, dominated by *Helicobacter*; shades of green are Bacteroidetes, dominated by *Bacteroides* and *Parabacteroides*; pinks and purples are Firmicutes, dominated by Lachnospiraceae and *Clostridium*; black represents unclassified bacteria.

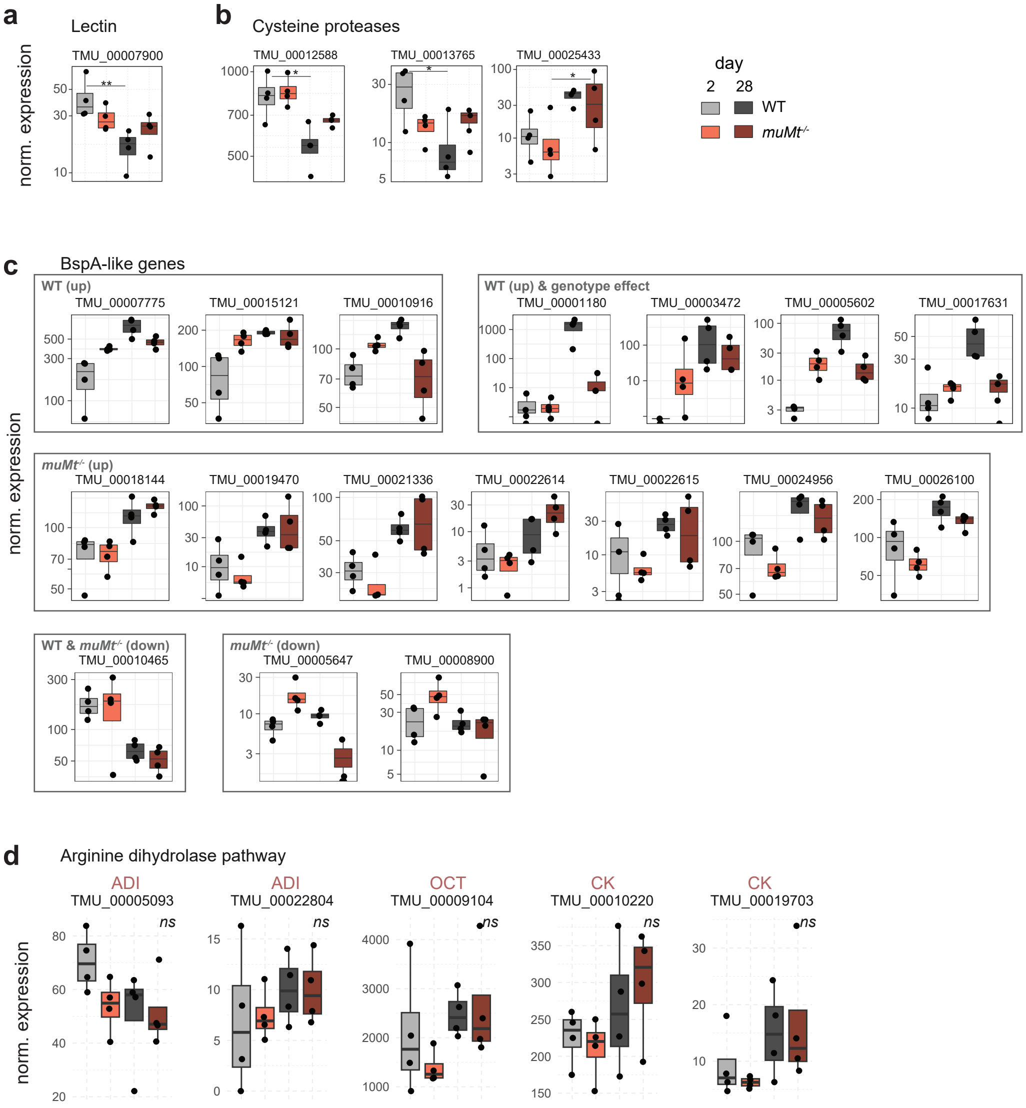


**Supplementary Figure 4.** Putative virulence-related *T. mu* genes differentially expressed over the course of colonization. **a**, Lectin and **b**, cysteine proteases differentially expressed at 28 compared to 2 days post colonization, in WT or *muMt*$^{-/-}$ mice as indicated. **c**, BspA-like genes with significant changes in expression over colonization time and/or which differ between host genotypes. **d**, Expression of genes with predicted activity in the arginine dihyrdolase pathway: arginine deiminase (ADI), ornithine carbamoyltransferase (OCT) and carbamate kinase (CK). Differences in gene expression were tested using DESeq2. *$p < 0.05$, **<0.01, ***<0.001, *ns* non-significant

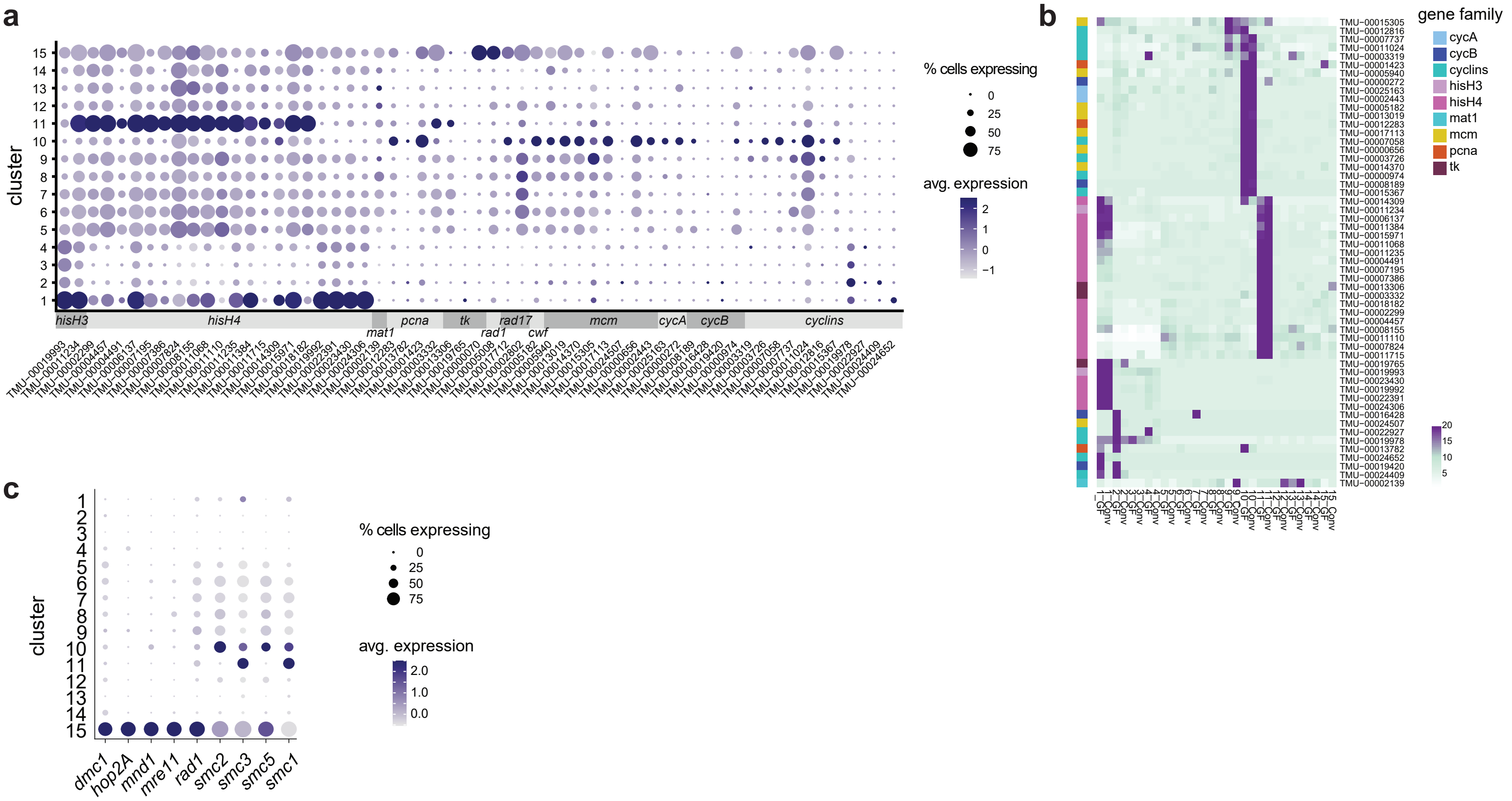


**Supplementary Figure 5.** Expression of cell cycle marker genes. **a**, Dotplot and **b**, heatmap depicting scaled read counts of genes known to be expressed during G1/S and G2 phases, across each *T. mu* cluster. Colour blocks in **b** (*left*) indicate assigned gene function. Clusters are separated by the host mouse. **c**, Expression of meiosis-specific genes.

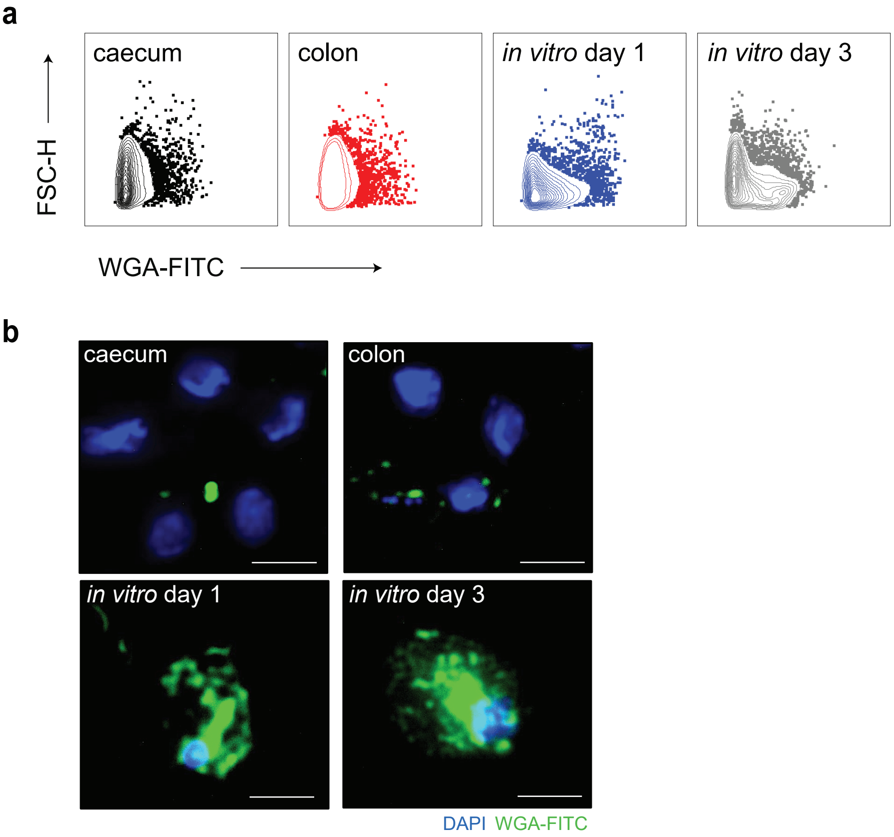


**Supplementary Figure 6.** WGA-staining of *T. mu* cells freshly isolated from mouse caeca or colons, or *in vitro* cultured for 1 to 3 days. **a,** Representative contour plots from FACS analysis of WGA-FITC stained protists. Events are gated on live, single protozoa. n=3 animals or culture plates per group, from four independent experiments. **b,** Cytospins of WGA-FITC (green) and DAPI (blue) stained protists. Representative images are shown at 63x magnification. Scale bars, 10 µm.

**a**

| | KEGG-defined pathway | ECs | ctl day 2 | ctl day 28 | *T. mu*-infected day 2 | *T. mu*-infected day 28 |
|---|---|---|---|---|---|---|
| Amino acid metabolism | Valine, leucine and isoleucine degradation | 7 | ns | ns | ns | * (3) |
| Carbohydrate metabolism | Glycolysis / Gluconeogenesis | 22 | ns | ns | ns | *** (7) |
| Lipid metabolism | Fatty acid degradation | 7 | ns | ns | ns | ** (4) |
| | **Total significant ECs** | | **433** | **166** | **349** | **349** |

**b**

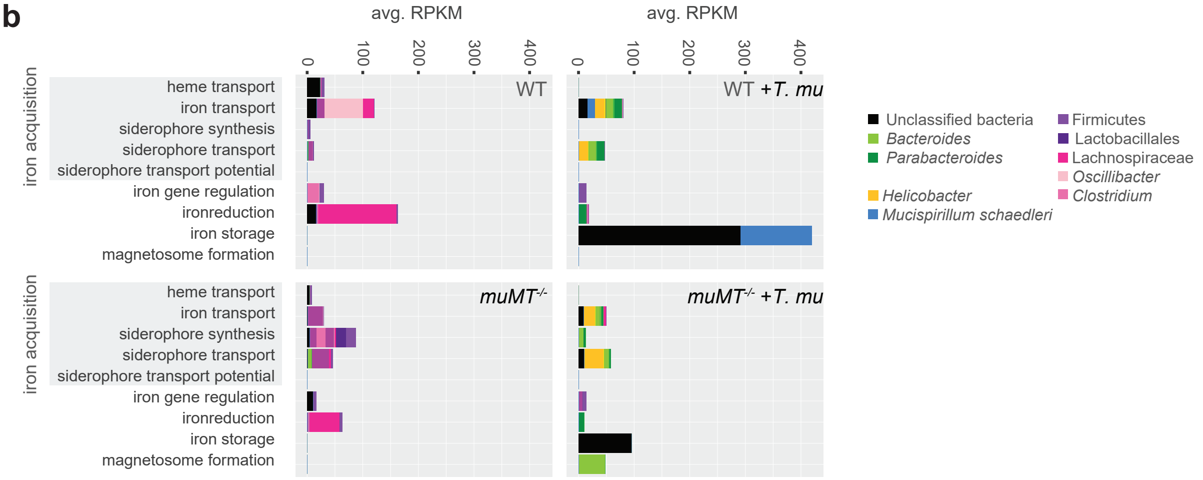


**c**

| iron acquisition: Fe category | *T. mu* infection day 2 WT | *T. mu* infection day 2 *muMT-/-* | *T. mu* infection day 28 WT | *T. mu* infection day 28 *muMT-/-* | host genotype (*muMT-/-* vs. WT) day 2 control | host genotype day 2 +*Tritrich.* | host genotype day 28 control | host genotype day 28 +*Tritrich.* |
|---|---|---|---|---|---|---|---|---|
| iron transport (630) | *ns* | *ns* | *ns* | *ns* | *ns* | *ns* | *ns* | *ns* |
| siderophore synthesis (799) | *ns* | *ns* | *ns* | *ns* | *ns* | *ns* | *ns* | *ns* |
| siderophore transport (1067) | *ns* | *ns* | *** (162) | *** (97) | *ns* | *ns* | *** (36) | *ns* |
| siderophore transport potential (134) | * (5) | ** (12) | ** (26) | *** (20) | *ns* | *ns* | *** (11) | *ns* |
| iron gene regulation (368) | *ns* | *ns* | ** (56) | *ns* | *ns* | *ns* | *** (19) | *ns* |
| iron reduction (50) | | *ns* | *ns* | *ns* | *ns* | | *ns* | |
| iron storage (33) | *ns* | *ns* | ** (10) | *ns* | *ns* | *ns* | | |
| magnetosome formation (23) | | *ns* | * (7) | ** (6) | *ns* | | *ns* | |
| | 642 | 1980 | 7199 | 4110 | 6753 | 5382 | 1075 | 337 |

**Supplementary Figure 7.** Functional enrichment of bacterial gene expression due to protist colonization and host genotype. **a**, Metabolic pathway enrichment in cecal bacteria, due to host genotype. **b**, RPKM of iron-related genes attributed to particular bacterial taxa at day 28 in WT or *muMT-/-* naïve and colonized mice. Colours represent taxa as indicated: black represents unclassified bacteria, greens are members of the Bacteroidetes phylum, pinks and purples are Firmicutes, yellows are Proteobacteria, blues are Deferribacteres. **c**, Enrichment of bacterial iron-related gene families due to protist colonization (*left*) and host genotype (*right*).
